# Synergistic computational and experimental studies of a phosphoglycosyl transferase membrane/ligand ensemble

**DOI:** 10.1101/2023.05.07.539694

**Authors:** Ayan Majumder, Nemanja Vuksanovic, Leah C. Ray, Hannah M. Bernstein, Karen N. Allen, Barbara Imperiali, John E. Straub

## Abstract

Complex glycans serve important functions in all living systems. Many of these intricate and byzantine biomolecules are assembled employing biosynthetic pathways wherein the constituent enzymes are membrane associated. A signature feature of the stepwise assembly processes is the essentiality of unusual linear long-chain polyprenol phosphate-linked substrates, such as undecaprenol phosphate in bacteria. In this study we focus on a small enzyme, PglC from <u>*Campylobacter*</u>, structurally characterized for the first time in 2018, as a detergent solubilized construct. PglC is a monotopic phosphoglycosyl transferase (PGT), that embodies the functional core structure of the entire enzyme superfamily and catalyzes the first membrane-committed step in a glycoprotein assembly pathway. The size of the enzyme is significant as it enables high level computation and relatively facile, for a membrane protein, experimental analysis. Our ensemble computational and experimental results reveal a specific interaction of undecaprenol phosphate with PGT cationic residues and suggest a role for critical conformational transitions and electrostatic steering in substrate recognition, overcoming significant energetic barriers to binding. The study highlights that computation, guided by fundamental chemical principles, can advance the study of biochemical processes at membrane bilayers and provide chemical insight at a molecular level that cannot be derived by experiment alone.

**Insert Table of Contents artwork here:** 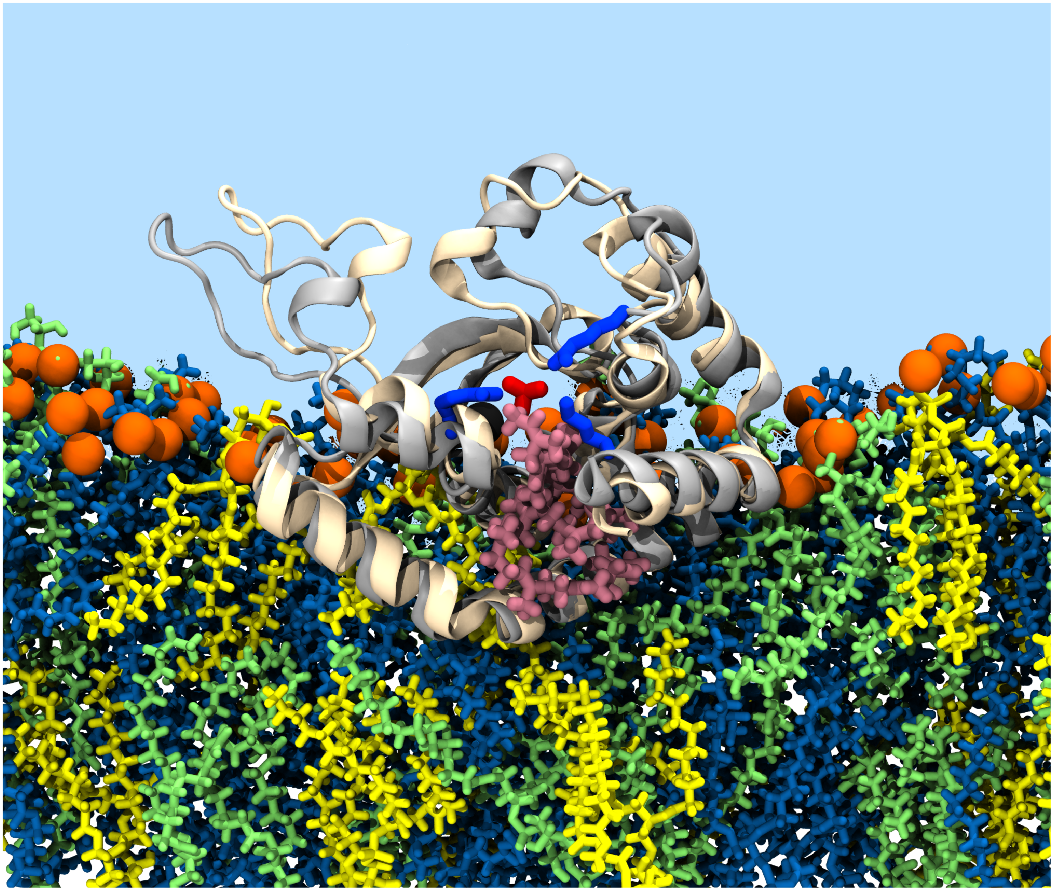

## Introduction

Understanding the dynamic interactions between membrane-bound enzymes and their membrane-associated substrates in a native lipid bilayer environment presents major logistical challenges. Currently, snapshot views from structural analysis, for example employing X-ray crystallography, can provide some insight into enzyme-substrate association. However, such analyses are most commonly conducted with detergent-solubilized proteins. This treatment inevitably disrupts native membrane interactions and precludes assessment of the contribution of the membrane environment to the subtle interplay of protein dynamics and ligand interactions that lead to function. The analysis of membrane-bound enzymes in their native environment has revealed unexpected properties that impact catalysis, as illustrated for example in the dramatic enhancement of catalysis observed for rhomboid proteases in membrane.^1^

Phosphoglycosyl transferases (PGTs) catalyze the transfer of a phosphosugar group from a UDP-sugar substrate to a membrane-resident polyprenol phosphate acceptor (in bacteria most commonly undecaprenol or decaprenol phosphate). ^2^ This reaction is the initiating step of many glycoconjugate biosynthesis pathways.^3-5^ The current studies focus on the monotopic PGT (monoPGT) superfamily.^4, 6^ The first structure for a monoPGT superfamily member, the *Campylobacter concisus* PglC (PDB 5W7L), was determined in 2018.^7^ This structure^7^ and extensive sequence analyses^6, 8^ show that all superfamily members include a minimal catalytic domain defined by a reentrant membrane helix (RMH) and an active site including an Asp-Glu catalytic dyad and magnesium cofactor positioned at the membrane interface. Subsequent computational analyses were also applied to define the sequence motifs that guide the RMH topology.^9^ Catalysis proceeds via a covalent intermediate^10^ with both soluble and amphiphilic substrates remaining in aqueous and membrane environments respectively.^11^ Although the novel structure provided new insights, the enzyme was crystallized in a detergent-solubilized form and lacked the native context of the membrane. Indeed, the snapshot of the structure that was captured represented an open conformation with direct access of bulk solvent to the active site. The suggestion of a catalytically competent closed conformer, which could lead to a protected active site, came from bioinformatic covariance analysis of PGT orthologs. The use of covarying residues to indicate structural contacts pointed to the interaction of the RMH with the catalytic core of the protein as well as the interaction of structural elements that would allow active site closure.^6, 8^ Additional information on the monoPGT and its membrane-bound substrate was also derived from studies using the model membrane styrene maleic acid lipoparticle (SMALP) method^12, 13^, which show that the amphiphilic UndP substrate is recruited to the PGT enzyme.^14^ Finally, in vivo analysis using the substituted cysteine accessibility method (SCAM) method^15^ and the observation of an ordered phosphatidyl ethanolamine head group in the crystal structure corroborated the monotopic topology and the placement of key membrane-interacting residues in PglC.^7, 9^

Major opportunities in the study of membrane proteins in native-like environments have emerged from the development of liponanoparticles such as Nanodiscs, stabilized by membrane scaffold proteins, introduced by Sligar and SMALPs, stabilized by amphiphilic copolymers such as styrene maleic acid (reviewed in ^16, 17^). Indeed, purified and stabilized samples of PglC using such membrane mimetic systems have been reported^13, 18^ However, although these systems are very attractive for some studies, certain limitations must be recognized. For example, the area of the membrane is limited (10-20 nm diameter), the boundaries between the membrane and the liponanoparticle scaffold are non-native and account for much of the available area, and additives such as divalent cations can be destabilizing. Importantly, the precision/resolution of experimental distance measurements such as FRET may be inadequate for the questions at hand.

Herein, we present a synergistic computational and experimental approach for analysis of dynamics of the monotopic PGT, PglC, and interactions with both the membrane and the unique membrane bound substrate undecaprenol phosphate. We apply the CHARMM36 force field, which is the most widely used all-atom resolution forcefield for lipid systems. This force field has been used extensively in simulations of eukaryotic, prokaryotic, and artificial membranes. Specifically, we validate a model representing the experimentally-determined catalytic core of PglC ca. 200 residues) and a lipid bilayer (400 phospholipids/200 per leaflet) of relevant composition and dimensions, with and without the amphiphilic substrate UndP. These studies show that the UndP adopts a compact conformation in a single leaflet of the bilayer; the inclusion of PglC shows that the UndP conformation mirrors the position of the reentrant helix making critical interactions with cationic residues. Molecular dynamics reveals motions that would promote a closed active site, which is also supported by the observation of conformers in an experimental crystal structure.

## Results and Discussion

### Simulated bacterial membrane exhibits a characteristic liquid-disordered phase

We first simulated a number of lipid bilayers without protein, to validate the membrane model against experimental observables including area per lipid. As a benchmark, we simulated a lipid bilayer composed of 67 mol% 1-Hexadec-anoyl-2-(9Z-Octadecenoyl)-sn-Glycero-3-Phosphoethanolamine (POPE), 23 mol% 1-Hexadecanoyl-2-(9Z-Octadecenoyl)-sn-Glycero-3-Phosphoglycerol (POPG), and 10 mol% cardiolipin (CL) of defined acyl-chain composition (**Figure S1**) to mimic the experimentally-determined average composition for typical inner membranes of Gram-negative bacteria including *Campylobacter concisus*.^19^ To define lipid organization and packing, the liquid crystal order parameter (P_2_) and 2D bond-orientational order parameter 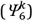 were computed for all lipid molecules. The P_2_ order parameter has been used to understand the organization of lipid tails in membrane domains, in both coarse-grained and all-atom models.^20, 21^ The value of the P_2_ order parameter reports on the orientation of the lipid director vector relative to the vector normal to the membrane. The range of P_2_ values varies from perpendicular (−0.50), to parallel (1.0). The P_2_ value for a liquid ordered domain of a lipid bilayer is greater than 0.9, using the CHARMM36 force field.^20^ In contrast, studies using the coarse-grained MARTINI model have reported P_2_ values varying from 0.63, for liquid disordered domains, to 0.82, for liquid ordered domains.^21^ The absolute value of the 2D bond-orientational order parameter 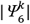 estimates the extent of hexagonal packing around the lipid. A high degree of hexagonal packing is a hallmark of the liquid ordered domain. The 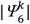 value varies from 0.42, for liquid disordered domains, to 0.48, for liquid ordered domains.^21^ The values of the P_2_ and 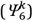 order parameters calculated from our simulations are tabulated in **Table 1**.

**Table 1:**
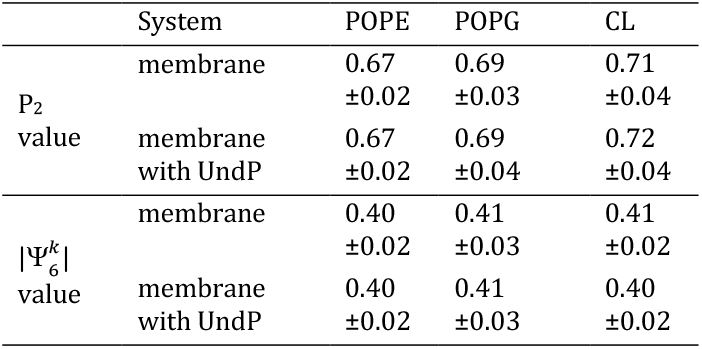
Values of P_2_ and 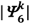 order parameters of different membrane components, measuring nematic order and hexagonal packing, respectively, obtained from simulation of membrane with and without the inclusion of UndP.

In comparison with previous studies, the lower values of the P_2_ and 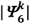 order parameters indicate that the lipid bilayer in this composition forms a liquid-disordered phase at room temperature. The distribution of the 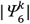 order parameter over different regions of the membrane shows no phase separation or domain formation in either leaflet (**Figure S2**). Parallel studies without CL showed that the major order parameters of the membrane components were not significantly perturbed (**Figure S3**).

### Undecaprenol phosphate (UndP) is disordered and localized in a single leaflet

Linear polyprenol phosphates (PrenPs) play a critical role in glycoconjugate biosynthesis and are present in concentrations estimated to be <0.1 mol% in bacterial membranes.^18, 22^ The structures of PrenPs have been explored through computational and experimental studies.^23-25^ Previous computational studies showed that dolichol (C95), dolichol phosphate (C95-P), and UndP feature three domains characterized by a central coiled region, involving a series of *Z*-configuration isoprene units, and two flanking regions, including the polar head group and the tail with *E*-configuration isoprene units.^24^ In a study of C95 and C95-P in a dimyristoylphosphatidylcholine (DMPC) bilayer, the polar head groups of the PrenP were observed to be located near the membrane-water interface with the polyprenyl coiled region interacting with the phospholipid acyl tails.

To investigate the behavior of UndP in this system, we placed two UndP molecules (1.0 mol%) in the lipid bilayer in an extended transmembrane conformation. The order parameters (Table 1) show that the inclusion of UndP does not perturb the lipid packing. This result is consistent with previous published studies employing pyrene as a probe to assess changes in membrane polarity, which showed only modest effects on the membrane up to 0.5% UndP.^14^ From the density analysis, the phosphoryl groups of the UndP molecules are observed to be primarily located near the membrane-water interface (**Figure 1b**). This density profile suggests that the UndP occupies the upper leaflet, in agreement with previous studies.^24^ We performed Voronoi tessellations by considering one atom per lipid tail to compute the area occupied by the different bilayer components (**Figure S4)**. The computed areas occupied by POPE, POPG, CL, and UndP were found to be 58.3, 57.7, 108.8, and 31.7 Å^2^, respectively. The average value of the radius of gyration of UndP, calculated over the course of the simulation (**Figure S4)**, was found to be 0.85 nm. Substantial fluctuations in the radius of gyration of UndP result from the flexibility of the polyprenyl group, which readily and frequently undergoes transitions between stretched, coiled, and unstructured conformations.

**Figure 1:**
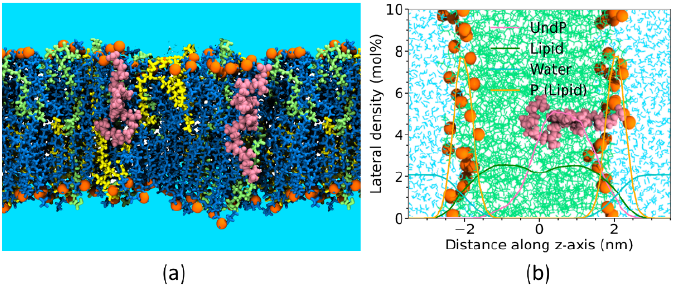
(a) Pictorial representation of the lipid bilayer composed of POPE (blue), POPG (green), CL (yellow), and UndP (salmon). Lipid head groups are represented in orange. (b) Density distribution of different membrane components. Traces represent the density distributions of UndP (salmon), lipid acyl chains (green), water (cyan), and phospholipid phosphate head group (orange), respectively.

### Validation of computational model of PglC in a model bacterial membrane

Having validated the membrane model in the absence and presence of UndP, we introduced the PglC protein employing the standard method for protein insertion using the CHARMM-GUI.^26^ The resultant placement was consistent with previous experimental and computational studies^7, 9^ (see Introduction). The experimentally determined structure of PglC (PDB 5W7L)^7^ was used to examine dynamics in the membrane environment (composed of 67 mol% POPE, 23 mol% POPG, and 10 mol% CL, as in the simulations above). We note the C-terminal residues (183-205) were not ordered in the X-ray crystal structure and therefore were excluded from the simulations in all the presented studies.

The RMSD values of the PglC structure over the time course, after the initial computational equilibration, with reference to the experimental structure, is shown in **Figure 2b**. The insertion depth of the protein obtained from the simulation is shown in **Figure S5**. The comparison of RMSF of PglC with the experimentally derived B-factors (**Figure S6**) shows excellent agreement between experiment and simulation. The X-ray crystallographic structure shows that the signature RMH (see Introduction) is kinked at the Ser23-Pro24 motif with an interhelix angle of 118°. In the simulation, the structure of the reentrant helix (residue 1-36) is similar to that in the crystal structure with a RMSD of < 1 Å. Those regions of PglC exposed to water show the largest structural fluctuations, although the average RMSD values over the simulation are less than 3 Å. The insertion depth of PglC observed in simulation is consistent with a reentrant helix with a hinge angle at Ser23-Pro24 of 117° consistent with experiment. The maximum insertion depth of PglC is 13 Å, also in agreement with structural and biochemical studies.^7, 9^ These findings validate our computational model of PglC in a model bacterial membrane.

**Figure 2:**
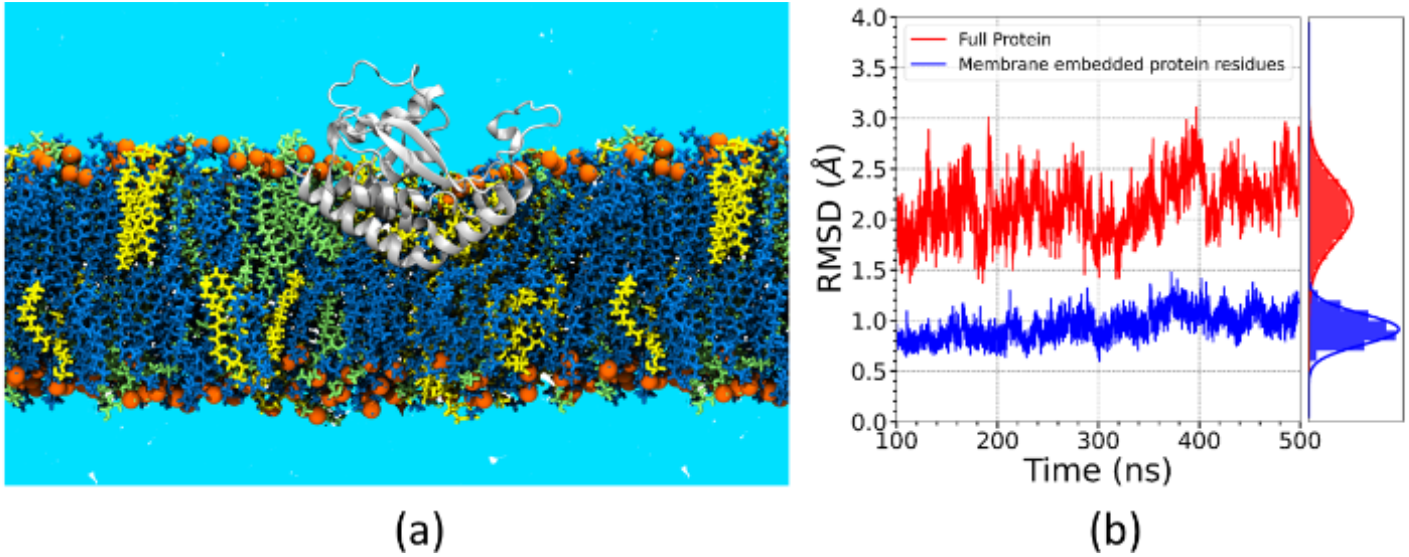
(a) Pictorial representation of the POPE:POPG:CL 67:23:10 lipid bilayer, showing POPE (blue), POPG (green), CL (yellow). The PglC protein and lipid head groups are represented by grey, and orange, respectively. (b) RMSD of PglC structures obtained from simulation, along with the cumulative distributions. The crystal structure was used as the reference state.

### Role of basic residues and electrostatic steering in UndP binding to PglC

Previously, we reported that the presence of PglC in a model membrane system increases the local UndP concentration by two-fold relative to the bulk membrane.^9^ It has also been shown that conserved basic residues in the active-site cavity of PglC play an important role in the binding of UndP to PglC.^8^ To identify key interactions between UndP and PglC, we simulated a lipid bilayer using the same composition and parameters as described above, augmented by the inclusion of two UndP molecules (0.5 mol %, see methods). This proportion of UndP is somewhat higher than a typical physiological concentration (<0.1 mol %) but is necessary to constrain the size of the system for computation. One UndP molecule was found to interact with PglC (**Figure 3a**). The second UndP did not show any specific binding and was observed to diffuse away from PglC over the course of the simulation. The contact map for PglC and the interacting UndP, obtained from simulation, is shown in **Figure 3b**. The resulting binding pose suggests that the interaction of UndP and PglC is mediated by Arg88, Arg145, and Lys179 (**Figure 3c**).

**Figure 3:**
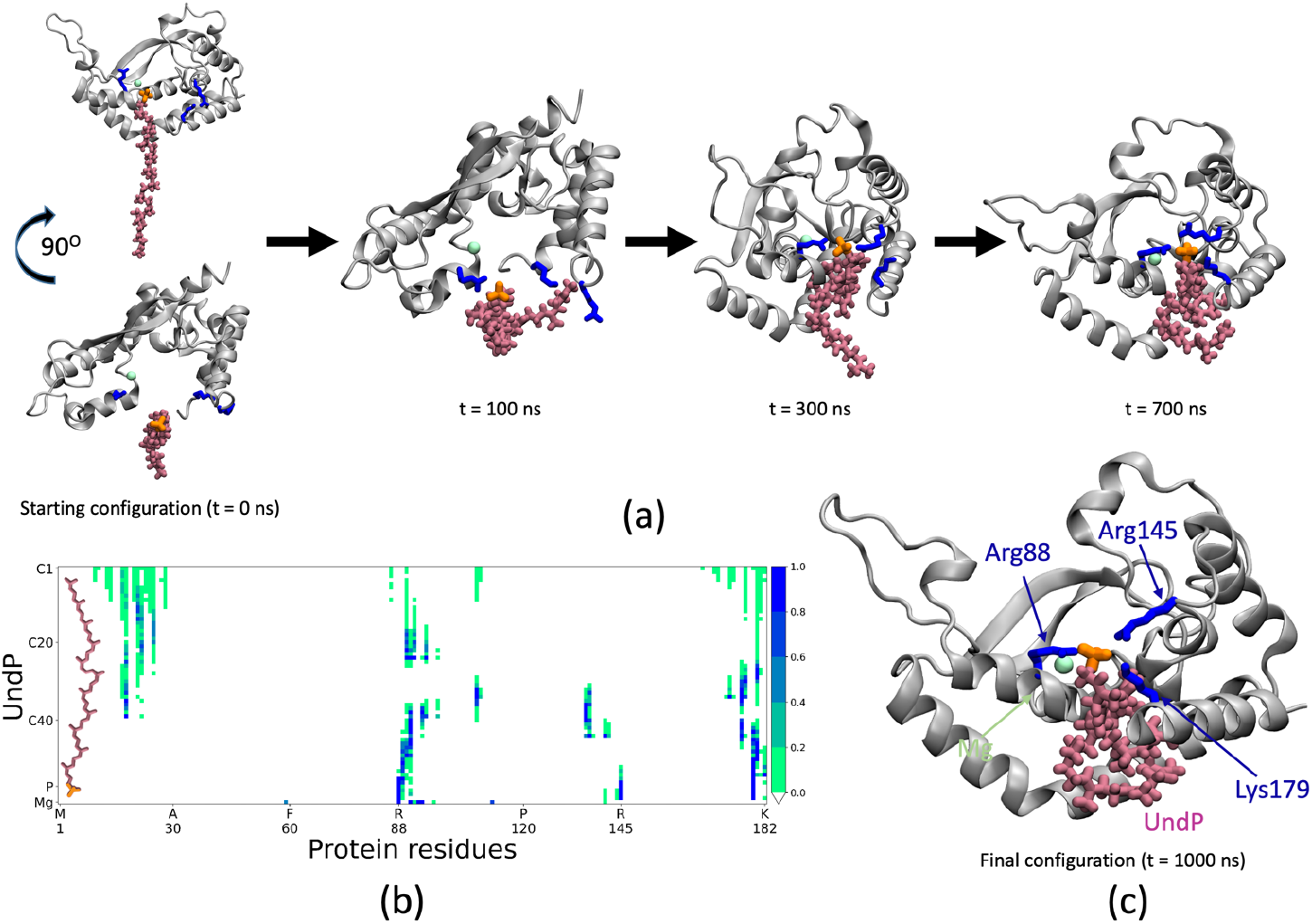
(a) Interaction of PglC and UndP at different times of the simulation. Lipids and water were not shown for clarity. (b) Contact map of UndP and PglC using a 5 Å interaction distance cutoff. The bar colors represent the contact probability, where a value of one represents a contact maintained between the residues throughout the simulation trajectory. (c) Ribbon representation of UndP and PglC interaction. Grey, salmon, green, and blue represent the PglC, UndP, Mg2+, and residues of PglC that interact with UndP, respectively. The location of the active site and catalytic DE motif is coincident with the Mg2+ cofactor.

The computational prediction of Arg88 as a mediator of UndP binding is consistent with a previous mutational analysis^6, 8^ showing that the R88Q variant has minimal activity (less than 30% wild type (WT) activity when tested at 10-fold higher concentration). In these studies, Arg145 has a large effect when substituted by either alanine or glutamine, with ∼100-200-fold reduction in activity when compared to WT for either variant. However, the mutation of Lys179 and Lys182 (singly or in a double mutant) to either alanine or glutamine indicated that these residues do not play a significant role in catalysis.

Considering these experimental results, we performed parallel simulations of three site-directed variants of PglC, R88Q, R145Q, and R88Q-R145Q, in a membrane with two UndP molecules, as described above. The UndPs were initially positioned as in the WT PglC simulation. In contrast to the simulation of the WT PglC, UndP was not observed to bind to the active site of any variant PglC during a 1 μs simulation. This observation is consistent with the experimental kinetic analysis of the variants; replacement of one or two basic arginine residues with a polar, but neutral, glutamine impacted the binding of the negatively charged UndP. To explore the role of charge in the relative binding affinity of UndP to the WT and variant forms of PglC, we computed the electrostatic potential map (**Figure 4**). In the case of WT PglC, the Arg88, Arg145, and Lys179 residues form a patch of positive charge near the active site (**Figure S7**). In contrast, the PglC variants have significantly diminished positive charge in the active site region. We postulate that this change in charge distribution disrupts the electrostatic steering of the UndP toward the active site and diminishes the binding affinity. In addition, the R88Q mutation deforms the region encompassing residues 174-182 of PglC (**Figure S8**). Similar patterns were seen in independent simulations of the R88Q and R88Q-R145Q variants. Along with the diminished positive charge density in the active site, the deformation of this (residues 174-182) region would be likely to negatively impact UndP binding to the active site of PglC.

**Figure 4:**
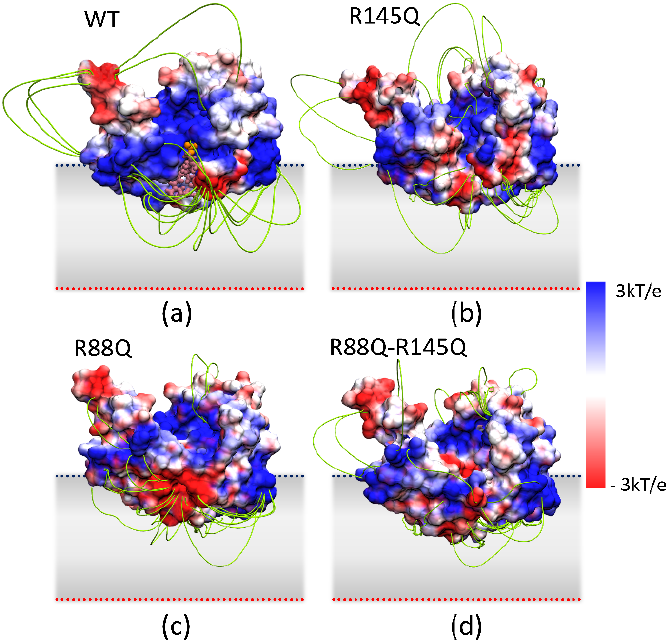
Electrostatic potential map of (a) WT, (b) R145Q, (c) R88Q, and (d) R88Q -R145Q PglC. The protein surface color, varying from red to white to blue, represents electrostatic potential values ranging from negative to positive values. The corresponding electric field lines are shown in green. UndP is depicted in the active site of WT PglC to indicate the position of the active site. UndP, and the head group of UndP are represented by salmon, and orange respectively. The grey layer represents the average position of the membrane.

### Closure of the mobile loop is correlated with occupancy of the active site

Closure of the mobile loop (*C. concisus* residues 61-81 see **Figure 5**), which connects the extended β-strand of the core fold of the monoPGT with the helix proximal to the active site, has been hypothesized to be integral to substrate binding.^7^ This model is based on two previous observations: (a) the divergence of the loop position between energy-minimized predicted structural models and the crystallographically determined model and (b) conservation of residues within the mobile loop. In the current study, further insight is derived from the structure of selenomethionine (SeMet)-derivatized PglC co-crystallized with UDP. In contrast to I57M/Q175M PglC,^7^ the SeMet PglC has an asymmetric unit (ASU) comprising eight monomers rather than two per unit cell, allowing independent refinement. Although the resolution was low (3.01 Å), and the ligand was not stoichiometrically bound (10 – 20 % occupancy of any ligand to the active site), it was possible to make correlations between the presence or absence of electron density and the loop conformations. When the eight copies of PglC in the ASU were overlaid, the mobile loop is the main region of structural divergence between the chains (**Figure 5a; Table S1**). In monomers G and H, which are closed, as well as A and B (intermediate conformation), a comparatively large number of electrons is present in the site where the nucleotide would be expected to bind. In contrast, monomers C, D, E, and F, which are open, all have significantly fewer electrons present (**Figure 5b**). Analysis of the mobile loop in the context of active-site ligand occupancy shows that chains with a greater positive electron density were in a more closed conformation than those with little or no density. Monomers occupying the same position within each of the four lattice dimers do not always occupy the same mobile loop position, thus the difference in the observed loop conformation is likely due to differential binding at the active site. Mapping of crystallographic B-factors in each monomer reveals that the loops occupy either one of two relatively low-B-factor conformations corresponding to open (C and D) or closed (G and H), with higher-B-factor loops occupying positions between these two extremes (A, B, E, and F) (**Figure 5c**).

**Figure 5.**
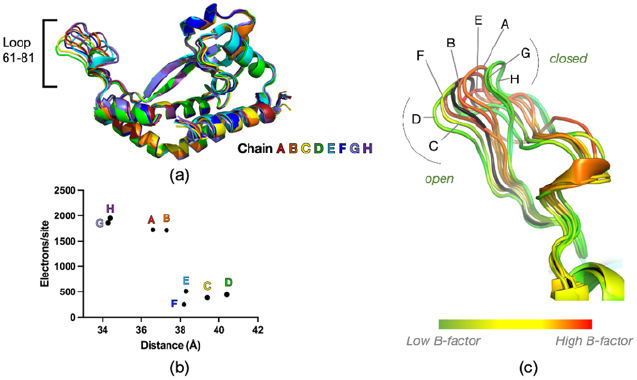
Mobile loop “closed” conformations observed upon occupancy of PglC active site. (a) Overlay of the 8 chains in the WT SeMet dataset. Chains colored by rainbow from chain A (red) to chain H (violet). (b) Estimate of the number of electrons in the active site of each chain derived using Mapman27 (see Methods). The distances were measured from the Cα at the top of the mobile loop (Cα of N70 in all chains except E, where it is Cα A69, due to differences in loop conformation) to the Cα of I121. (c) B-factors for each residue mapped to the mobile loop. Low B-factors are observed for the “open” and “closed” conformations, correlated with electron density in the active site. For reference, the structure of I57M/Q175M PglC at 2.74 Å resolution (PDB 5W7L) is shown in black.

### Correlation of mobile loop dynamics in simulation and experiment

We investigated the loop-closing motion of PglC using two independent means: structures derived from our computer simulations and a series of eight experimentally-derived PglC crystal structures. In both cases we apply principal component analysis (PCA) to identify essential motions of the protein.^28, 29^ The resulting PCs provide insight into conformational differences of the protein in the crystal environment and observed in molecular dynamics simulations (**Figure 6a** and **b**).

**Figure 6:**
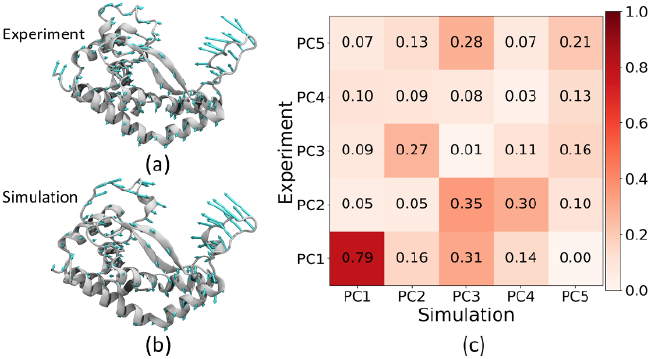
Movement of the protein along first principal component obtained from (a) experiment and (b) simulation. (c) Inner product of the eigenvectors associated with each PC obtained from experiment and simulation.

In both cases, we found that the loop closing motion corresponds to the eigenvector characterizing the first principal component. To provide a quantitative comparison of the two predicted principal components, we projected one principal component onto the other to determine the quantitative overlap or similarity. To compare the PCs from structures derived from experiment and simulation, we computed the inner product of the first five PCs. The inner product varies from highest (1, identical PCs) to lowest (0, no similarity) degree of overlap ((**Figure 6c**). In each case, the first PC is dominated by motion of the mobile loop of PglC. We observe a high degree of similarity for the first PCs, demonstrating strong correlation between the large-scale loop dynamics observed in simulation, in the membrane environment, and obtained from experiment, in the crystal environment.

Our results qualitatively and quantitatively demonstrate the similarity between the loop motion in experiment and simulation. This synergistic comparison provides insight into the primary functionally important motion in PglC.

## Conclusions

The challenge of how to investigate both dynamics and interactions within a membrane/ligand/protein ensemble was addressed herein using computation augmented by experiment. We have adopted an iterative approach using computation to yield a functionally relevant view of a monoPGT in the membrane and used the results both to interpret and inform experimental design. The all-atom molecular dynamics simulation, using a lipid bilayer composition representative of the inner membrane of Gram-negative bacteria, was observed to form a liquid-disordered phase at room temperature, a property which was maintained upon addition of UndP. In this ensemble, the phosphate headgroups of the UndP are found at the membrane-water interface, whereas the acyl tails were disordered and largely localized within a single leaflet of the model bacterial membrane. Such an arrangement is expected to be enthalpically equivalent to an extended transmembrane conformation but entropically favored.

Assessment of the monoPGT in the simulation showed good agreement between the experimentally determined X-ray crystal structure (solved with detergent-solubilized protein) and the equilibrated computational structure in the membrane. The PglC structure, which is substantively embedded in the membrane (24% embedded in contrast to the 3.9% average for monotopic membrane proteins)^7^, was found to undergo fluctuations with low RMSD values (< 3 Å) in comparison to the crystal structure. The N-terminus of the protein forms a reentrant membrane helix by creating a 117° angle at the hinge centered on the Ser23-Pro24 residues. These computationally derived observations of the protein within the lipid bilayer correlated well with observations derived from experiment.

The analysis also provided a high-level view of the membrane embedded PglC/UndP ensemble. The findings further suggested that it is advantageous for the polyprenol phosphate to largely occupy the leaflet where the reentrant membrane helix resides as opposed to additionally disrupting the opposing leaflet of the bilayer, which would be energetically costly. The computational approach yielded the first view of a polyprenol phosphate binding to a monoPGT active site, revealing that the basic residues Arg88, Arg145, and Lys179 of PglC play a critical role in facilitating the recognition and binding of UndP. This observation highlights a specific mode of interaction of UndP with enzyme and is consistent with experimental results showing the activity of PglC was significantly reduced when Arg145 or Arg88 were mutated to Ala or Gln. The importance of these residues is underscored by the computed electrostatic potential and associated electric field, which suggests a role for electrostatic steering. Such electrostatic steering likely acts as a major driving force, contributing to recognition and binding of both UndP and the soluble nucleotide sugar substrate. Both substrates must overcome significant energetic barriers in binding to the active site − desolvation in the case of the UDP-sugar and competition with lipid headgroup interactions in the bulk membrane in the case of UndP.

The application of theory elucidated the molecular basis underlying experimental observations and validated differences in structural conformations as functionally relevant. It was previously conjectured that closure of the mobile loop may play an important role in substrate binding and attainment of the catalytically competent conformation. Our analysis shows excellent agreement between the simulated ensemble of protein structures in membrane and the experimentally observed structures with varying degrees of active site-ligand occupancy in the crystal environment. In both cases, we observe global fluctuation of PglC between open and closed states, characterized by the position of the mobile loop relative to the active site. Taken together, our results provide a holistic view of the critical role of charge and structural transitions in the recognition and binding of the UndP substrate to a monoPGT and highlight the synergistic use of computation and experiment to assess membrane-resident interactions. More generally, in the case of membrane proteins where structure determination is so challenging and biophysical characterization in model membrane systems has limitations – approaches integrating high-level simulations can help to advance understanding of this critical sector of the proteome.

## Methods

### Computational Models and Analysis Methods

PglC and UndP were simulated in a lipid bilayer composed of 67 mol% POPE, 23 mol% POPG, and 10 mol% CL of defined acyl-chain composition (**Figure S1**) using the CHARMM36m all-atom force field.^30, 31^ A lipid bilayer composed of 200 phospholipid molecules without PglC was prepared. A second lipid bilayer composed of 400 phospholipid molecules containing UndP and PglC (PDB ID 5W7L)^7^ was also prepared. Residues 57 and 175 of PglC were mutated to Ile and Gln, respectively, using Chimera^32^ as these mutations were present in the experimental protein structural analysis (and constitute the WT protein used in this study). Each lipid bilayer was solvated using the TIP3P water model.^33^ The concentration of KCl was set at 0.15 M. The UndP-only and UndP-plus-PglC systems were solvated with 52 and 80 water molecules per phospholipid, respectively. Each bilayer was equilibrated for a minimum of 50 ns of molecular dynamics simulation in the constant pressure and temperature ensemble, using the Nose-Hoover thermostat and Parrinello-Rahman barostat, following the CHARMMGUI protocol.^26^ A 1.5 μs production run was performed for the UndP-only system and a 1 μs production run was performed for the systems containing UndP and PglC. The temperature of each production run was maintained at 303 K. All simulations were performed using the GROMACS 2018.3 program.^34^

The liquid-crystal order parameter (P_2_) was computed for lipids using the angle (θ) between the director vector defined by a subset of phospholipid atoms and the bilayer normal vector where:

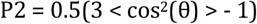

For POPE and POPG, the director vectors were defined by the C1 through C16 carbon atoms and C1 through C14 carbon atoms. For CL, the director vectors were defined by the C1 through C12 carbon atoms (**Figure S1**).

The 2D bond-orientational order parameter 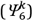 was calculated by considering one atomic coordinate per lipid chain using:

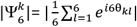

where θ_*kl*_ is the angle between an arbitrary vector and the vector connecting the central atom (*k*) and one of the six nearest-neighbor atoms (*l*). Voronoi tessellation was also performed considering the position of the lipid chains. For POPE and POPG, the coordinates of the C14 and C16 carbon atoms, and for CL, the coordinates of the C12 carbon atoms, were used to define the positions of the lipid chains (**Figure S1**). The electrostatic potential maps were calculated using PDB2PQR^35^,APBS^36^, and APBSmem.^37^ Principal component analysis for the protein was performed by diagonalizing the covariance matrix of the position of the backbone atoms using ProDy^38^ and GROMACS tools. All other analyses were performed using in-house code written using the MDAnalysis Python library.^39^

### Preparation of Site-Directed Variants

Single and double point mutations were introduced into the WT *C. concisus* SUMO-PglC sequence in the pE-SUMO vector^7^ using the QuikChange II Site-Directed Mutagenesis kit (Agilent) and primers in **Table S2**.

### Protein Purification

WT and SUMO-PglC variants were expressed and purified using published protocols.^7^ Briefly, proteins were expressed in *E. coli* BL21-DE3-RIL cells by autoinduction. Cell pellets were resuspended in base buffer (50 mM HEPES pH 7.5, 100 mM NaCl) supplemented with 25 mg lysozyme, 25 μL DNAse I and 50 μL protease inhibitor cocktail. Cells were sonicated twice for 1.5 minutes (1 second on/2 seconds off, 50% amplitude), and the lysed cells were centrifuged at 9417 g for 45 minutes. The resulting supernatant was centrifuged at 142,414 g for 65 minutes to pellet the cell envelope fraction (CEF). The CEF was homogenized into base buffer with 1% DDM and rotated overnight at 4°C. The detergent-homogenized sample was centrifuged at 161,571 g for 65 minutes. The supernatant was incubated with 1 mL Ni-NTA resin for 1 hour at 4°C. The resin was washed with 20 column volumes of Wash I Buffer (base buffer, 0.03% DDM, 20 mM imidazole, 5% glycerol), followed by 20 column volumes Wash II Buffer (base buffer, 0.03% DDM, 45 mM imidazole, 5% glycerol). SUMO-PglC was eluted in 2 column volumes of Elution Buffer (base buffer, 0.03% DDM, 500 mM imidazole, 5% glycerol) and immediately desalted using a 5 mL HiTrap Desalting Column into 50 mM HEPES pH 7.5, 100 mM NaCl, 0.03% DDM, 5% glycerol. Protein purity was assessed by SDS-PAGE with Coomassie staining (**Figure S9**).

### Activity Assays

Assays for PGT activity were performed using the UMP-Glo assay (Promega). Reactions contained 20 μM UndP, 20 μM UDP-diNAcBac, 50 mM HEPES pH 7.5, 100 mM NaCl, 5 mM MgCl2, 0.1% Triton X-100, and 10% DMSO with 0.4 nM SUMO-PglC. Reactions in the linear range were quenched with the UMP-Glo detection reagent and luminescence was measured in a plate reader as previously described.^9^ Relative luminescence units were converted into concentration of UMP (μM) with a standard curve.

### Crystallization

WT-SeMet *C. concisus* PglC used for crystallography was prepared as described previously.^7^ Notably, PglC co-purifies with approximately 3 phosphatidyl ethanolamine : 1 phosphatidyl glycerol endogenous lipid observed by thin layer chromatography. WT Se-Met PglC crystals were grown by hanging drop vapor diffusion at 17 °C from a 1:1 mix of protein and well comprised of 0.1 M Bis-Tris pH 6.0, 0.3 M MgCl_2_, 27% PEG 3350, and 1 mM TCEP (2 μL total volume). Se-Met PglC (276 μM in in 50 mM HEPES pH 7.5, 100 mM NaCl, and 0.03% DDM) was used after incubation with 1 mM UDP on ice for 30 minutes. WT-SeMet PglC crystals used for data collection appeared within 3 days and reached their final size after 14 days. Crystals were flash-cooled by plunging into liquid nitrogen for transport and data collection without additional cryo-protection.

### Data Collection and Refinement

The WT-SeMet PglC dataset was collected at BNL NSLS-II 17-ID-1 (AMX) (Upton, NY) at the Se X-ray absorption energy peak (12665 eV) allowed initial partial phases to be solved by SAD using the Phenix suite.^40^ Matthews coefficient analyses for the data set was consistent with 8 copies in the asymmetric unit. Data were scaled and integrated using XDS.^41^ SHELXD^42^ was run for 5000 trials with a resolution cut-off of 4.5 A to identify 16 Se sites. Phenix.SOLVE^43^ was used to find an additional 6 Se sites and calculate subsequent Se substructure phases for 22 out of the expected 32 Se atoms in the ASU. Phenix.RESOLVE^44^ was used to perform initial solvent flattening and phase-extension. The partial model resulting from these maps was utilized, together with data from a second native I57M/I87M 2.59 Å dataset to determine the published structure of the more complete, higher I/σ(I) dataset of I57M/Q175M PglC at 2.74 Å resolution (PDB 5W7L).^7^ This higher resolution model was ultimately used to calculate the phases for the WT-SeMet PglC dataset.

Refinement against the electron density map was performed with Phenix.Refine^45^ to refine XYZ coordinates, realspace, rigid body, and group B-factors. Subsequent rounds of refinement included refinement of translation libration-screw (TLS) parameters, manually placed waters, and simulated annealing of Cartesian coordinates and torsion angles. The final model with eight subunits in the asymmetric unit was refined to R_work_/R_free_ of 0.27/0.30 with no significant outliers using Phenix.Refine^45^. All chains of the model contain 185 out of 205 amino acids as density corresponding to the C-terminus was not visible. Data collection and refinement statistics are tabulated in **Table S2**. The coordinates were deposited in the PDB under entry 8E37.

### Electron Density Analysis

The mFo-DFc map of WT-SeMet *C. concisus* PglC in CCP4 format was generated from an MTZ file containing coefficients using PHENIX^40^. The electron density in the active site of PglC protomers was quantified using MAPMAN^27^. The peaks in mFo-DFc map at 3.0 σ were identified and the electron density in a 3.5 Å sphere surrounding peaks was integrated to quantify the number of electrons in the volume where UDP is expected to bind^46^.

## Supporting information

ASSOCIATED CONTENT

## ASSOCIATED CONTENT

(Table S1) mutagenesis primers; (Table S2) data collection and refinement statistics; (Figure S1) lipid structures; (Figure S2) lateral lipid organization; (Figure S3) area of lipids in the absence of CL (Figure S4) area of lipids and radius of gyration of UndP; (Figure S5) insertion depth of PglC; (Figure S6) comparison of RMSF and B-factor; (Figure S7) position of basic residues in WT and variant form of PglC; (Figure S8) overlay of WT and variant forms of PglC; (Figure S9) SDS-PAGE analysis.

## ACKNOWLEDGMENT

The authors gratefully acknowledge the generous support of the National Institutes of Health, grant R01 GM131627 (to K.N.A. and B.I.) grant R01 GM107703 (to J.E.S.), National Science Foundation, grant no. CHE1900416 (to J.E.S.), and the high-performance computing resources of the Boston University Shared Computing Cluster (SCC).

## Notes

### Competing Interest Statement

The authors have declared no competing interest.

