## Supplementary material for "Synergistic computational and experimental studies of a phosphoglycosyl transferase membrane/ligand ensemble": ASSOCIATED CONTENT

**Table S1: *C. concisus* PglC variant site-directed mutagenesis primers**

| construct | primer | |
|---|---|---|
| R145A | primer #1 | CGCAGGTAAATGGCGCAAACGCCATAAGTTGGG |
| | primer #2 | CCCAACTTATGGCGTTTGCGCCATTTACCTGCG |
| R145Q | primer #1 | CGCAGGTAAATGGCCAAAACGCCATAAGTTGGGAG |
| | primer #2 | CTCCCAACTTATGGCGTTTTGGCCATTTACCTGCG |
| K179A | primer #1 | GCCTTACAGACAATAGAAGCGGTGCTAAAACGAAGTGG |
| | primer #2 | CCACTTCGTTTTAGCACCGCTTCTATTGTCTGTAAGGC |
| K179Q | primer #1 | GCCTTACAGACAATAGAACAGGTGCTAAAACGAAGTG |
| | primer #2 | CACTTCGTTTTAGCACCTGTTCTATTGTCTGTAAGGC |
| K182A | primer #1 | CAATAGAAAAGGTGCTAGCACGAAGTGGTGTCAGCAAAG |
| | primer #2 | CTTTGCTGACACCACTTCGTGCTAGCACCTTTTCTATTG |
| K182Q | primer #1 | CAATAGAAAAGGTGCTACAACGAAGTGGTGTCAGC |
| | primer #2 | GCTGACACCACTTCGTTGTAGCACCTTTTCTATTG |
| K179A/ K182A | primer #1 | CCTTACAGACAATAGAAGCGGTGCTAGCACGAAGTGGTGTCAGC |
| | primer #2 | GCTGACACCACTTCGTGCTAGCACCGCTTCTATTGTCTGTAAGG |
| K179Q/ K182Q | primer #1 | CTTACAGACAATAGAACAGGTGCTACAACGAAGTGGTGTC |
| | primer #2 | GACACCACTTCGTTGTAGCACCTGTTCTATTGTCTGTAAG |

**Table S2. Data collection and refinement statistics**

| | WT SeMet PglC PDB ID 8E37 |
|---|---|
| **Data collection** | |
| Beamline | BNL NSLS-II 17-ID-1 (AMX) |
| Wavelength (Å) | 1.0<br>0 |
| Resolution range (Å) | 47.737 – 3.013 (3.121- 3.01) |
| Space group | P 31 2 1 |
| Unit Cell (Å) | a = b = 142.82,<br>c = 192.563 |
| Total Reflections | 75337 |
| Unique reflections | 38779 (3544) |
| Multiplicity | 1.9 (2.0) |
| Completeness (%) | 82.76 (76.47) |
| Mean I/σ(I) | 14.2 (1.0) |
| Wilson B-factor | 83.8 |
| $R_{merge}$ | 0.074 (0.656) |
| $R_{meas}$ | 0.105 (0.928) |
| $CC_{1/2}$ | 0.98 (0.553) |
| **Refinement** | |
| Reflections used in refinement | 38391 (3428) |
| Reflections used for $R_{free}$ | 1994 (183) |
| $R_{work}$ | 0.2657 (0.3539) |
| $R_{free}$ | 0.2959 (0.3532) |
| Number of non-hydrogen atoms | 12088 |
| Protein residues | 1480 |
| RMS(bonds) | 0.002 |
| RMS(angles) | 0.61 |
| Ramachandran favored (%) | 95.90 |
| Ramachandran outliers (%) | 0.34 |
| Rotomer outliers (%) | 0.00 |
| Clashscore | 7.73 |
| Average B-factor | 111.53 |
| Number of TLS groups | 8 |

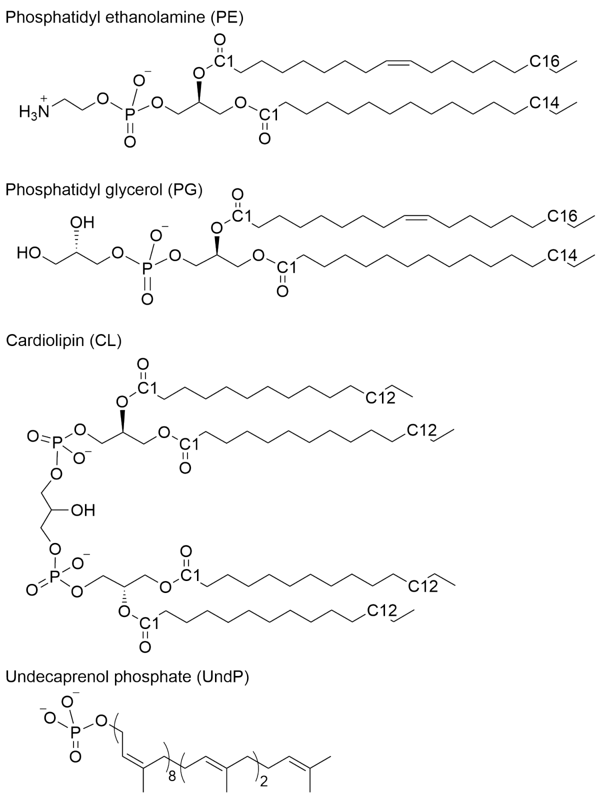


**Figure S1:** Structures of lipids included in the model bacterial membrane used in the simulation. The most common acyl chain modifications were used for each lipid. C1, C12, C14, and C16 atom positions of the lipids obtained from the trajectory were used for analysis.

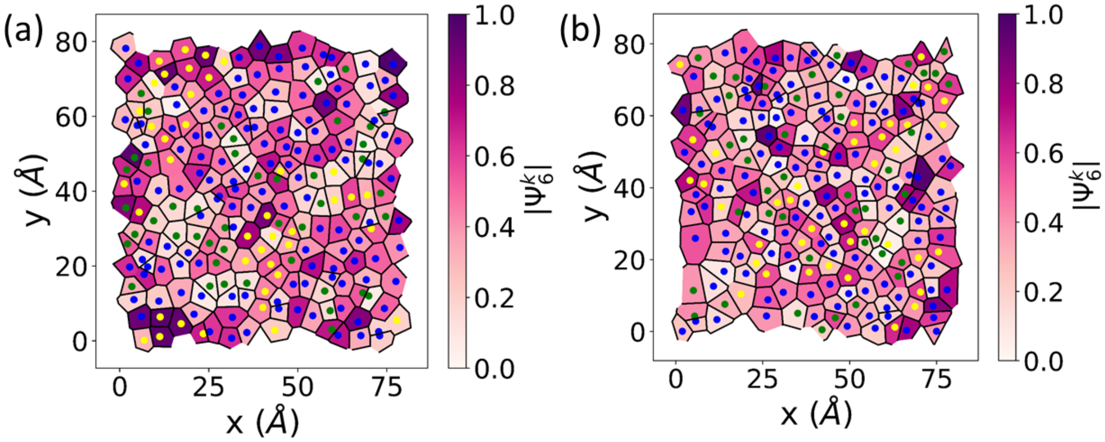


**Figure S2:** Voronoi tessellation showing lateral organization of lipids in the (a) upper and (b) lower leaflets in the final frame of the simulation. POPE, POPG, and CL are represented by blue, green, and yellow dots in center, respectively. Color of cell represents the value of the $|\Psi_6^k|$ order parameter providing a measure of local hexagonal packing.

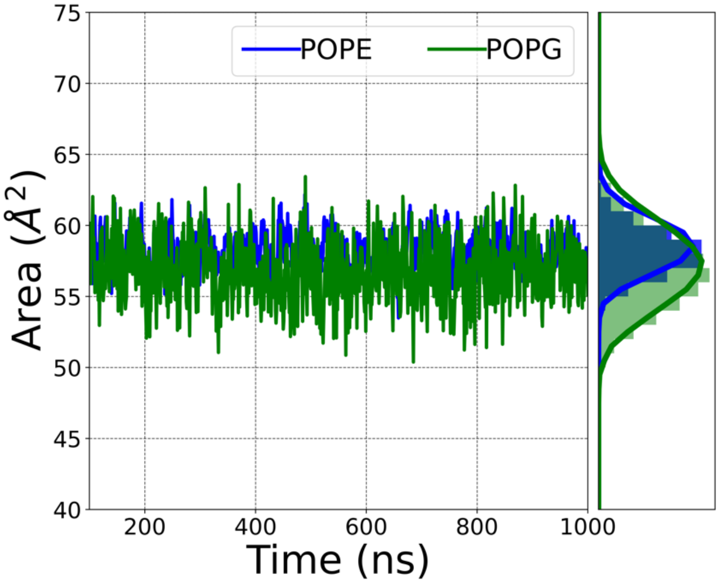


**Figure S3:** Instantaneous area of lipid components obtained by performing a Voronoi tessellation shown as a time series over the course of the simulation as an aggregate distribution for POPE (blue), and POPG (green). The simulation was performed with a membrane bilayer of composition 74 mol% POPE and 26 mol% POPG (no CL).

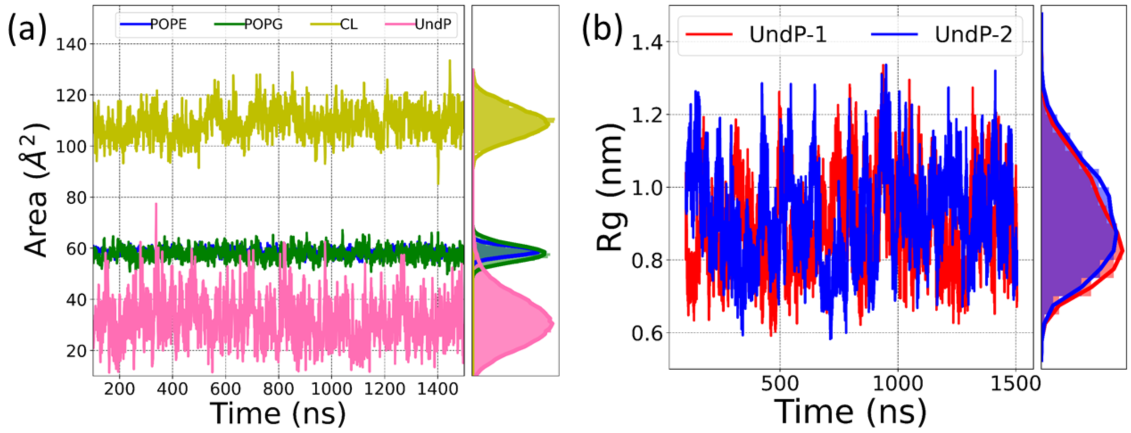


**Figure S4:** (a) Instantaneous area of lipid components obtained by performing a Voronoi tessellation shown as a time series over the course of the simulation an as aggregate distribution for POPE (blue), POPG (green), CL (yellow), and UndP (mauve). (b) Radius of gyration of UndP molecules shown as time series and aggregate distribution.

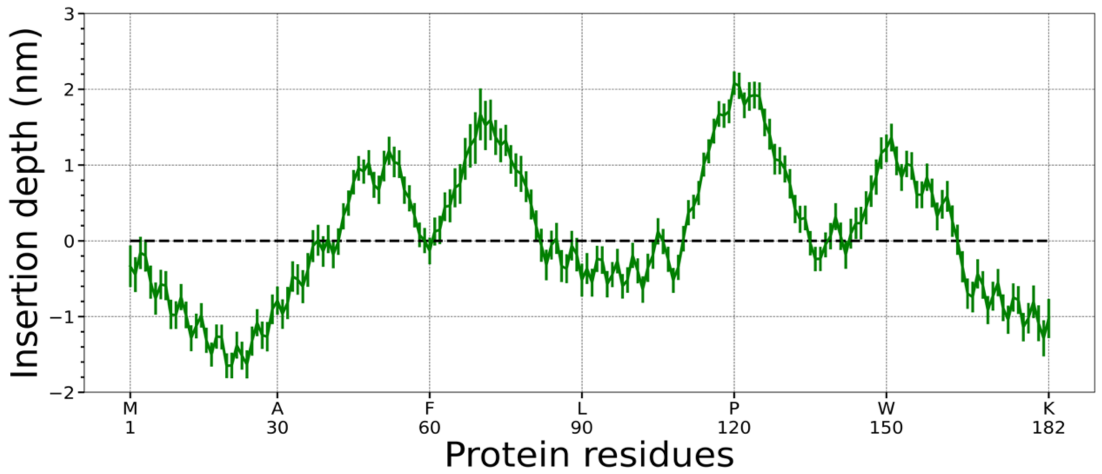


**Figure S5:** Average depth of insertion of PglC residues in the membrane bilayer averaged over the simulation. Oscillations indicate presence of helices. Error bars represent one standard deviation in the distribution of insertion depth.

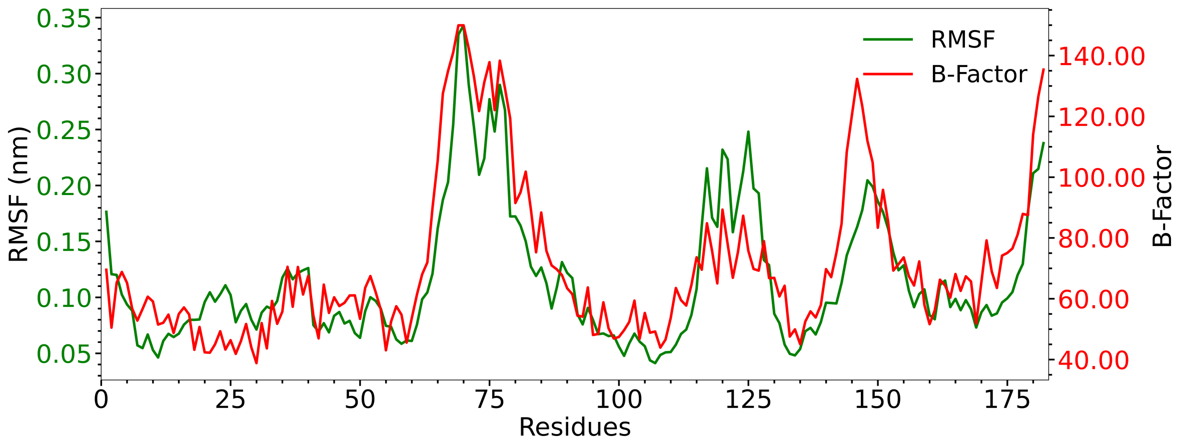


**Figure S6:** RMSF of PglC structures obtained from simulation and B-factor values obtained from the experiments.

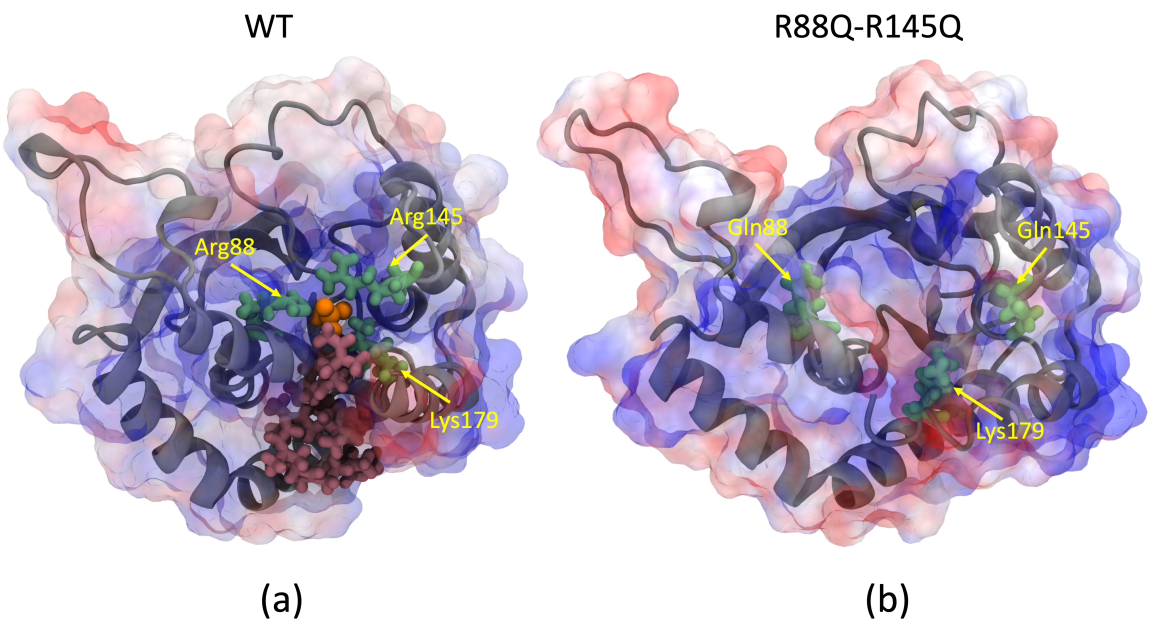


**Figure S7:** Ribbon diagram with overlayed electrostatic potential surface of (a) WT and (b) R88Q-R145Q PglC showing difference in the proximity of basic residues near the active site.

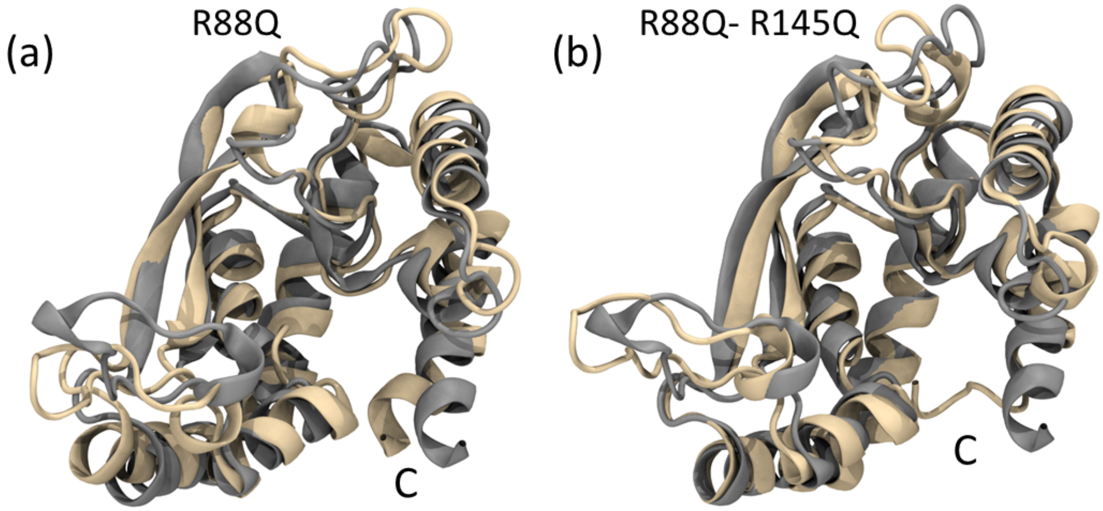


**Figure S8:** Overlay of representative instantaneous structures of (a) WT (gray) and R88Q PglC (wheat) and (b) WT (gray) and R88Q-R145Q PglC (wheat). The C-terminus of the PglC is marked by "C".

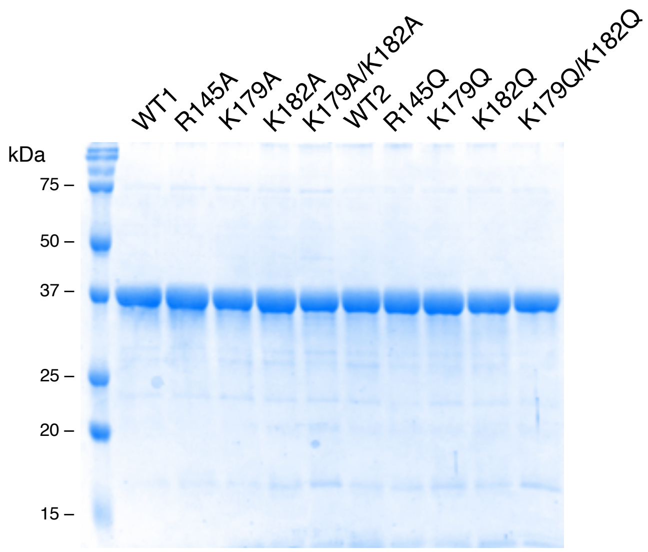


**Figure S9:** SDS-PAGE analysis of purified His$_6$-SUMO-PglC proteins. His$_6$-SUMO-PglC WT (technical duplicates WT1 and WT2) and variants made by site-directed mutagenesis (label across top of gel) were obtained employing Ni-NTA affinity chromatography. The expected molecular weight of the tagged proteins is 35.8 kDa.
